# Maternal patterns of inheritance alter transcript expression in eggs

**DOI:** 10.1101/2022.10.10.511534

**Authors:** Nathan D. Harry, Christina Zakas

## Abstract

Modifications to early development can lead to evolutionary diversification. The early stages of development are under maternal control, as mothers produce eggs loaded with nutrients, proteins and mRNAs that direct early embryogenesis. Maternally provided mRNAs are the only expressed genes in initial stages of development and are known to be tightly regulated. Differences in maternal mRNA provisioning could lead to phenotypic changes in embryogenesis and ultimately evolutionary changes in development. However, the extent to which variation in maternal mRNA provisioning impacts ontogeny or life-history is unknown. Here, we use a species with dimorphic development— where females make eggs and larvae of different sizes and life-history modes—to investigate the extent of variation in maternal mRNA provisioning to the egg. We examine the effect of gene expression differences on subsequent generations of egg provisioning and determine the regulatory architecture underlying mRNA provisioning differences. We find that there is significant variation in gene expression across eggs of different development modes, and that both parent-of-origin and allele-specific effects contribute to mRNA expression differences. We also find that offspring of intraspecific crosses differentially provision their eggs based on their parents’ cross direction. This effect of allelic expression based on parent-of-origin has not been previously demonstrated in reproductive traits like oogenesis.

**AUTHOR SUMMARY:** Variation in early developmental programs can provide the basis for evolutionary diversification. In the early embryo, cellular functions are carried out by proteins and transcripts contributed by the mother to the egg until the embryo’s own genome can take over. Since these maternal factors are responsible for setting up all of the subsequent development of the offspring, they tend to be tightly regulated. However, variation exists in the amount and types of transcripts mothers provide. Here we examine how the variation in maternal transcripts that occurs in eggs of the species *Streblospio benedicti*, leads to developmental differences. *S. benedicti* offspring follow one of two distinct developmental programs that originate with egg size differences. We find significant variation in maternally provided transcripts correlated with the two life-histories, and that some of this variation in egg transcripts is directly related to the developmental type of the mother’s own parents. This parental effect on how mothers provide transcripts to their eggs has not previously been described and indicates the possibility for an offspring’s grandparents to affect their early developmental program by modulating the transcripts their mother provides.

## INTRODUCTION

The unfertilized egg is a key time point to understand how differences in maternal provisioning affect ontogeny. In all metazoans, the first stages of embryonic development are carried out entirely by maternal proteins and mRNAs loaded into the oocyte. Control of development only transfers to the zygotic genome after a few cell divisions during a period called the maternal-to-zygotic transition, when the zygotic genome is activated by maternal transcription factors^(1–5)^. Because maternal transcripts directly control the offspring’s initial development, variation in maternal mRNA composition and abundance could profoundly alter subsequent developmental processes^(6–9)^. Changes in maternal mRNA expression may allow for shifts in development that ultimately lead to changes in life-history traits^(10–14)^.

How do differences in egg size and provisioning alter development? Variation in egg size, even in offspring with the same zygotic genotypes, can alter life-history traits^(15,16)^. Egg size is a cornerstone trait of life-history theory as well as developmental biology, but while egg size is often used as a proxy for maternal investment, size and provisioning do not always scale^(17–19)^. Maternal RNA provisioning may be variable with respect to egg size. For example, across cichlid species egg size is linked with differences in maternal RNA deposition, especially in growth related genes^(20)^. In frogs, maternal RNA localization differs in species with different egg sizes^(21,22)^ and RNA content changes with naturally occurring egg size differences within species^(23)^. When egg size is linked to life-history mode, it is unclear whether transcript abundance will change with egg size so that larger eggs receive a different amount of mRNA (quantitative change), or if completely different mRNAs are provided in conjunction with metabolic demands or new cellular functions (qualitative change).

### Study System

To investigate how mRNAs are provisioned to eggs of different sizes (and consequent development modes), we use the marine annelid *Streblospio benedicti*. This model has a developmental dimorphism: there are two types of females found in natural populations that produce eggs of different sizes: 100μm and 200μm diameter eggs that have an eight-fold difference in volume. The offspring develop into different larval types with alternate life-histories categorized by their trophic mode. Small eggs develop into planktotrophic larvae (obligately plankton feeding) and large eggs develop into lecithotrophic larvae (yolk-feeding^(24,25)^).

Planktotrophic mothers produce hundreds of small eggs per clutch and the larvae develop a gut and swimming structures early in development^(26,27)^. By comparison, lecithotrophic mothers produce tens of larvae per clutch with no swimming structures. Lecithotrophic larvae have an abbreviated larval phase, different larval morphologies, and require only maternally provided energy to undergo metamorphosis^(28)^. Despite these developmental differences, both larval morphs converge on the same body plan after metamorphosis. This intraspecific developmental dimorphism provides an opportunity to study how development can evolve from differences in egg provisioning.

Previous work in *S. benedicti* shows the genetic basis of alternative larval phenotypes is modular, with independent loci affecting individual traits^(29)^. For egg size in particular, there are both parental and zygotic loci that affect size^(16)^. However, the parental loci can act in both paternal and maternal effect directions^(30)^. This means that egg size is partially determined by inherited alleles, but also by the genotype of its parents. The mechanism by which these parental effects shape development and egg size is unknown. Here we explore if mRNA expression differences in the oocytes can account for developmental differences due to parental genetic effects.

As a single species, adults that arise from the two larval types of *S. benedicti* can be crossed to produce viable F_1_ offspring. Crosses can be reciprocal, alternating parental roles between a planktotrophic and a lecithotrophic adult. F_1_ females produce eggs of intermediate size compared to their parents, which will develop into F_2_ offspring with segregating larval traits^(16,29)^ (Figure 1). This allows us to use F_1_ females’ eggs to disentangle the effects of egg size and mRNA expression.

**Figure 1.**
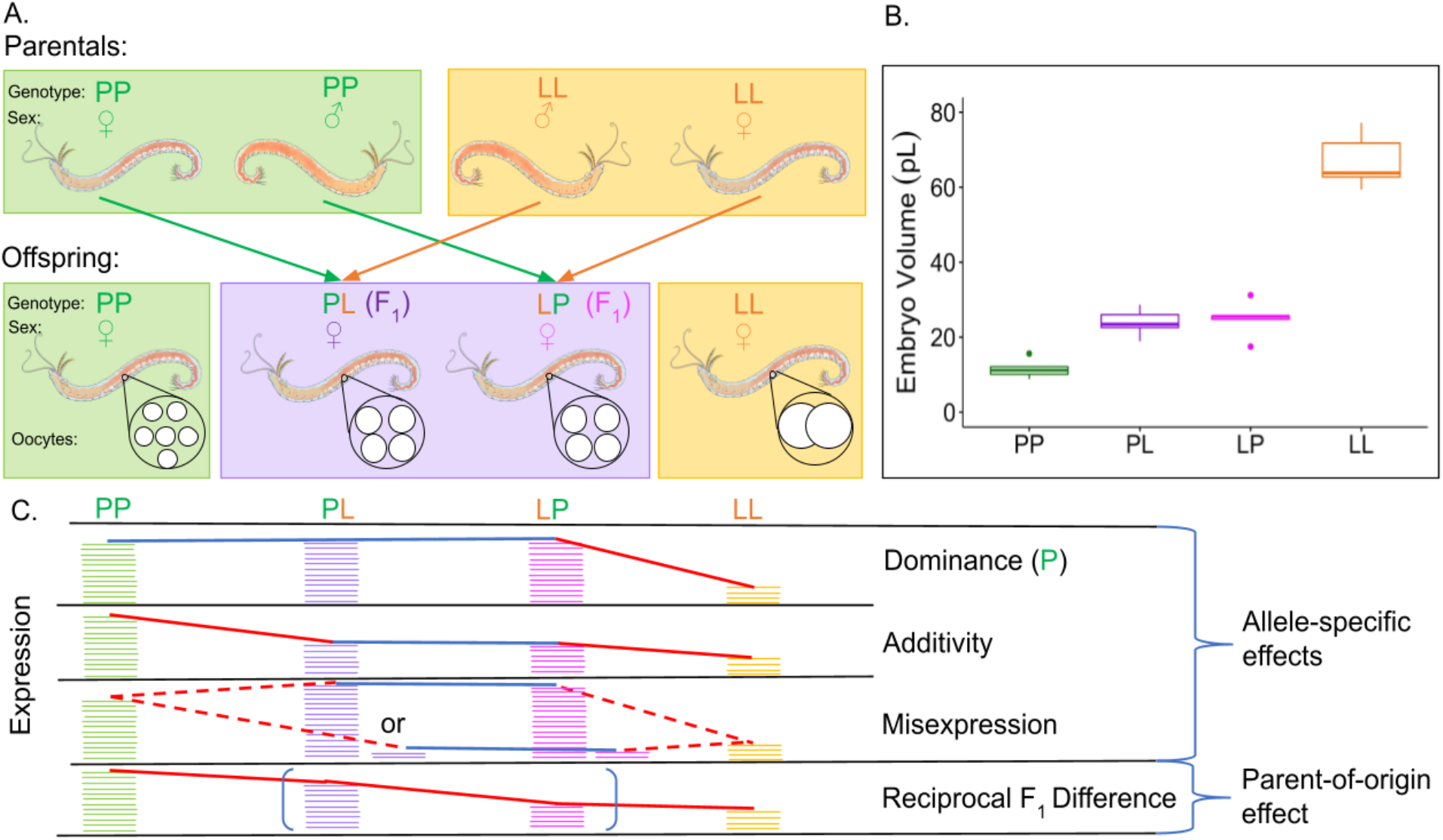
(A) Top: Reciprocal cross schematic to generate F_1_ females. Bottom: Representative females and eggs generated from crosses. (B) Early embryo area as a proxy for egg size in the four categories of females used in this study. F_1_ females have intermediate egg sizes compared to the parental types. Number of clutches measured for P = 6, L = 5, PL = 8, LP = 4 (10 embryos measured per clutch). (C) Allele-specific expression is when parental strains have differential expression, and PL and LP have matching expression levels relative to each other. A parent-of-origin effect is when the reciprocal F_1_s have differing expression levels relative to each other and the parentals. Blue lines indicate the same expression levels and red lines indicate a decrease in expression. Misexpression can be either over or underdominance.

### Approach

In this study we leverage the differences in egg size and development of *S. benedicti* to understand how maternally provided transcripts are associated with evolutionary developmental changes. We determine whether the evolutionary transition between planktotrophy and lecithotrophy is correlated with a qualitative or quantitative change in mRNA provisioning by comparing gene expression in the two types of eggs.

We use the eggs of F_1_ females from crosses between the two developmental types to investigate the regulatory architecture of mRNA expression changes. Using F_1_ females’ eggs allows us to disentangle **allele-specific effects** on expression from **parent-of origin effects** (Fig. 1C). We determine the mode of inheritance for differentially expressed genes and show the extent of allelic dominance and additivity. In contrast to allele-specific expression, we also determine the effects of parent-of-origin: where the expression level depends on if the allele originated from the mother or father. Parent-of-origin differences in gene expression between reciprocal F_1_s indicate an intergenerational effect on oocyte mRNA provisioning; isolating allele-specific and parent-of-origin inheritance patterns allows us to understand the extent to which these mechanisms contribute to variation in mRNA expression in eggs. However, this analysis is only possible in species where reciprocal crosses are viable, which is rare when considering crosses of individuals with dramatically different egg sizes and developmental modes. In fact, no studies to date have demonstrated an intergenerational effect on egg mRNA provisioning.

By using eggs of F_1_ mothers we can determine the regulatory architecture of differentially expressed genes: expression differences can be attributed to *cis* or *trans*-acting genetic factors (reviewed in: ^(31)^). Allele-specific differences in hybrids or strain-crosses have been used to investigated regulatory changes in a number of models: yeast^(32–34)^, flies^(35–39)^, plants^(40,41)^, fishes^(42,43)^, and mice^(44,45)^. Such studies have demonstrated that gene regulatory differences between species can be predominantly driven by either *cis* or *trans*-acting factors, and this varies depending on the species crossed. Studies in *Drosophila* that account for evolutionary divergence times across species show that *cis*-acting factors are greater in interspecific hybrids than for intraspecific F_1_s^(38)^. We test for the effects of *cis* and *trans* regulatory modifications on differentially expressed genes in the eggs made by F_1_ mothers (the eggs that would give rise to the F_2_ generation). Therefore, we can determine if mRNA provisioning differences are due to *trans or cis*-acting regulatory elements in the maternal genome. As *S. benedicti* is a single species with little genome-wide differentiation^(46,29)^, this analysis shows how genetic divergence of coupled reproductive and life-history transitions may begin to evolve.

## RESULTS

### Comparison of LL and PP egg gene expression

Reciprocal crosses generated F_1_ females from P and L parents (Fig. 1) and principal component analysis (PCA) indicate that we capture significant biological differences between groups, as the samples separate by developmental mode on the first principal component (Fig. 2). We found 1,155 genes that are differentially expressed in oocytes of the two developmental modes. These genes make up 10.24% of total expressed genes (n = 11,161) and 4.4% of the total number of genes in the *S. benedicti* reference genome (n = 26,216). 515 genes (44.6% of the DEGs) are expressed more in planktotrophic oocytes, and 640 are expressed more in the lecithotrophic oocytes (Fig 2). Of those 1,155 differentially provisioned genes, 92 (8%) are non-coding RNAs.

**Figure 2.**
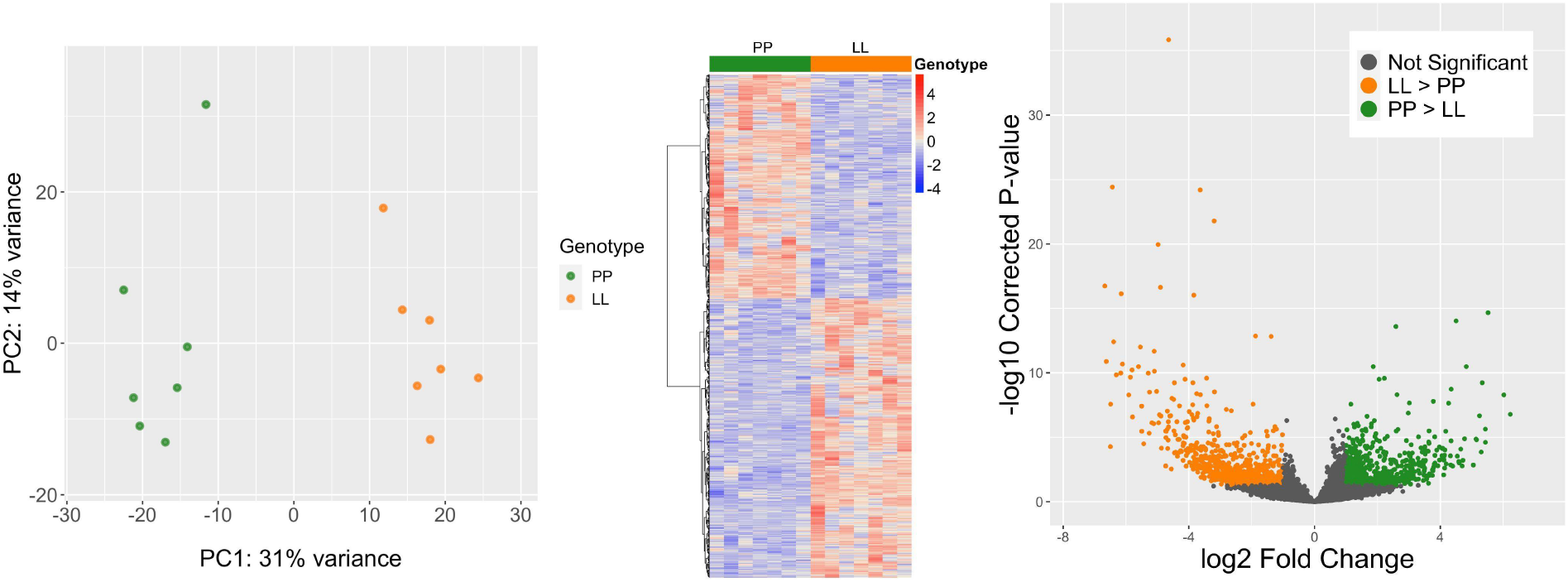
Differential Gene expression between planktotrophic and lecithotrophic oocytes. (A) PCA (B) Heatmap (C) Volcano plot

Genes that are only expressed in one of the morphs are considered exclusive. 18 genes have very low (sample group mean read count is fewer than 10 reads) or no expression in one morph. The planktotrophic eggs express eight exclusive genes and the lecithotrophic eggs express ten exclusive genes. Most of these genes are unknown in function (STable 5).

### Comparison of reciprocal F_1_ gene expression with LL and PP

When the F_1_’s eggs are included in the PCA (Fig. 3a) we found they fall between the parentals on PC2, but the variance among F_1_ samples is high (Fig. 3a,b). Most transcripts (78%, 8,722 genes) have conserved expression levels among all eggs. Out of the 1,155 genes that are differentially expressed between PP and LL eggs, 504 of the genes could be confidently assigned a mode of inheritance based on expression in F_1_’s eggs (Fig. 4a,b). 429 genes are dominant for one morph, 4 are additive, and 71 are over or underdominant. The remaining 651 DEGs either were not sufficiently sequenced in the F_1_ samples (uninformative) or have an ambiguous expression pattern likely due to high variation with F_1_ oocyte samples. The number of genes exhibiting dominant expression is almost even between P and L-allelic dominance, only slightly skewed in the L direction.

**Figure 3.**
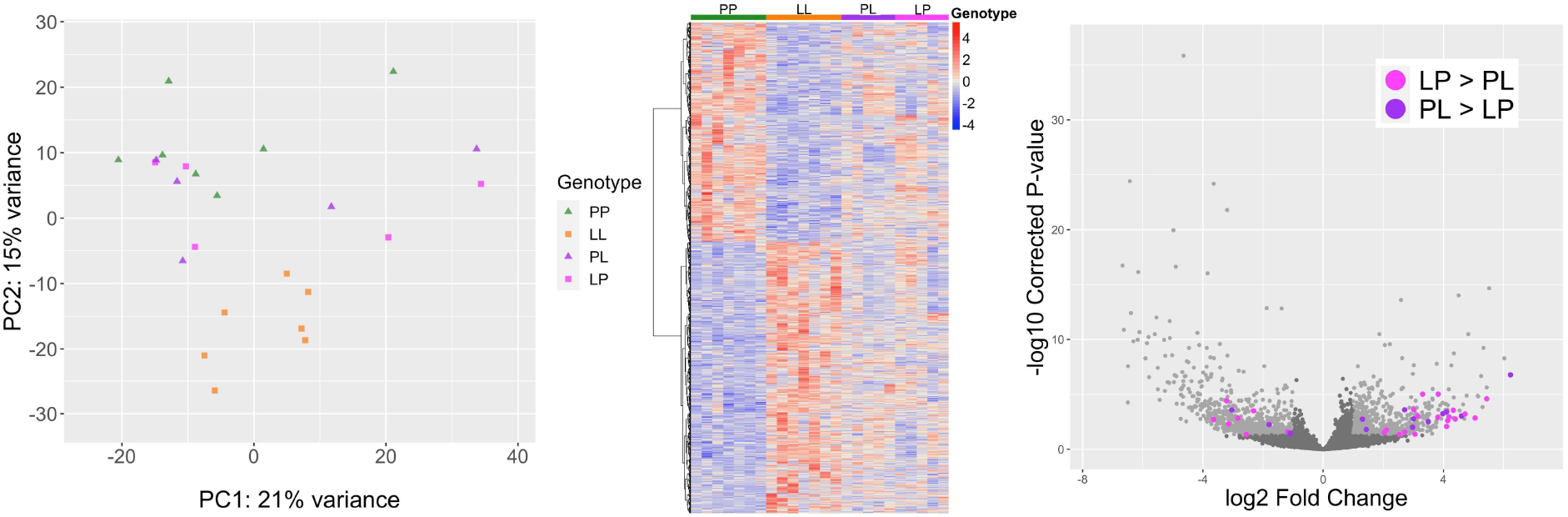
Differential Gene expression between PL and LP oocytes. (A) PCA shows F_1_ expression is intermediate to the parentals. (B) Heatmap shows both directions of F_1_’s gene expression alongside PP and LL expression (C). Volcano plot of differential expression between P and L with 38 genes that are also differentially expressed between F_1_ crosses highlighted. Color corresponds to the direction of expression change between the F_1_ crosses.

**Figure 4.**
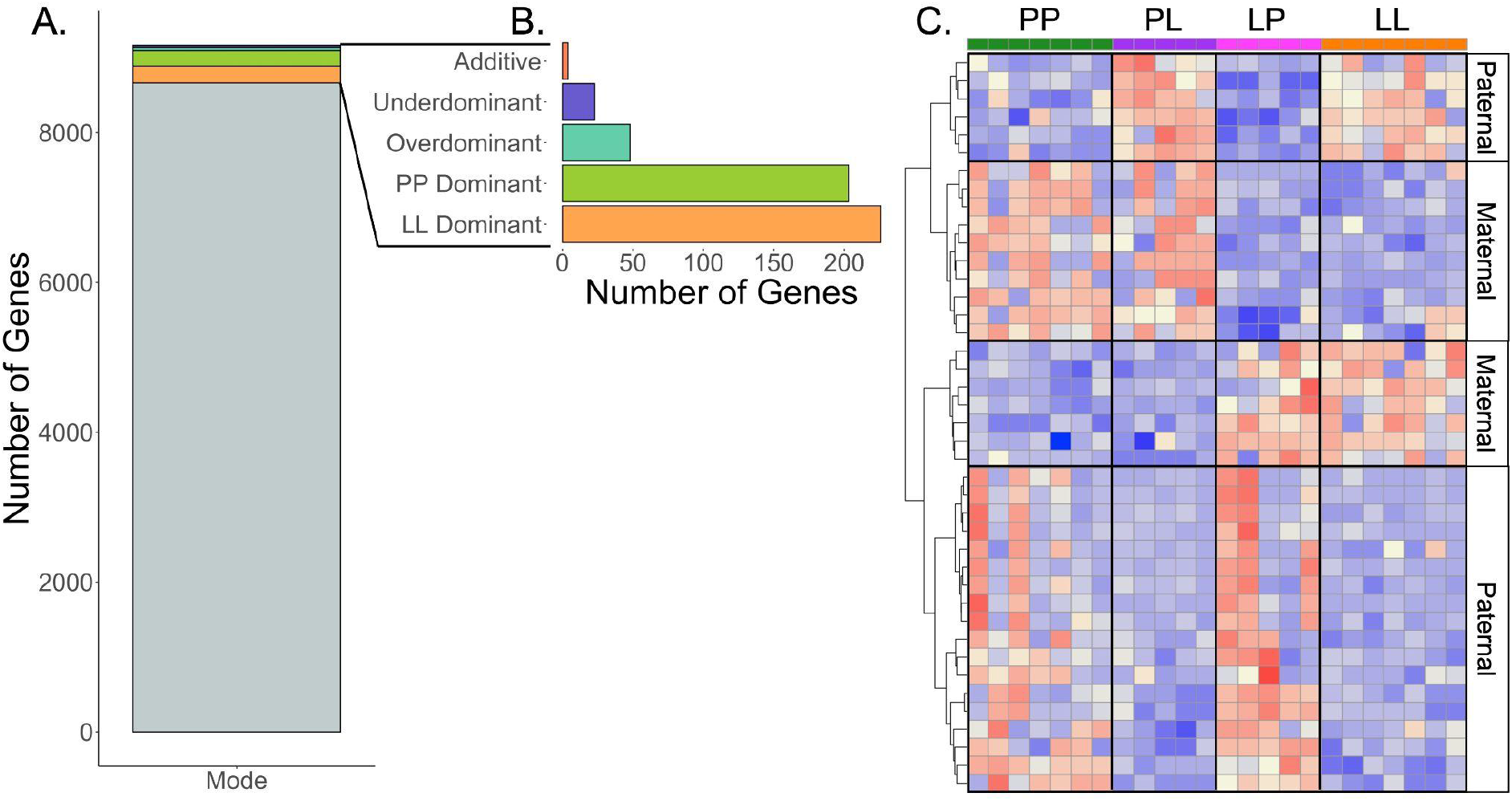
Mode of inheritance underlying expression divergence in eggs within a single species. (A) 8,722 genes are categorized as “conserved” (no expression level difference among parental or offspring’s eggs). (B) 523 of the remaining 1,155 genes that are differentially expressed between PP and LL eggs have been classified by their primary mode of inheritance. 102 genes are found to be differentially expressed between PL and LP. (C) A heatmap of genes with statistical support for parental effect direction. The expression direction (red is high expression and blue is low expression) for these genes matches the direction of either parental type. For example, if PP and PL have the same direction of expression (same color) this is a maternal effect, whereas if PL matches the direction of LL it is a paternal effect on expression. 24 genes are classified as primarily paternally inherited, and 17 as maternally inherited.

We compared the gene expression of the oocytes produced by reciprocal F_1_s (PL versus LP) to determine to what extent parent-of-origin effects influence maternally-loaded transcripts. Both F_1_ females are heterozygotes with respect to P and L alleles, so the only difference between PL and LP expression is the direction of the cross that gave rise to the mother (distinguishing if the allele is inherited from the maternal or paternal side). Genes that are differentially expressed between LP and PL are expressed in either the maternal or paternal direction, matching the expression of the PP parent or the LL parent. 102 genes are significantly differentially expressed between PL and LP (Fig. 3c), which accounts for 1.58% of the total number of genes expressed in oocytes. By comparing the expression of these genes back to PP and LL parental genotypes, we are able to infer that 24 are exhibiting paternal effects, and 17 are exhibiting maternal effects (Fig. 4c,d)

### Mode of regulatory change

Many differentially expressed genes could be co-regulated by a few *trans*-acting factors in the maternal genome. To test this possibility, we used SNP differences in the transcripts of F_1_’s eggs to assign each read to either the P or L allele. This analysis was only able to assign parental origin to 21.6% of F_1_ reads as there are few differentiating SNPs in this intraspecific comparison. We assigned the regulatory mode for 96 genes. We found that genes whose expression could be explained by *cis*-regulatory differences between alleles was only slightly greater than the genes with *trans*-acting regulation (Fig. 5). There are only a few *cis* + *trans* or of *cis* x *trans* changes, and some genes exhibit compensatory expression patterns (n=19; Fig. 5, STable 3).

**Figure 5.**
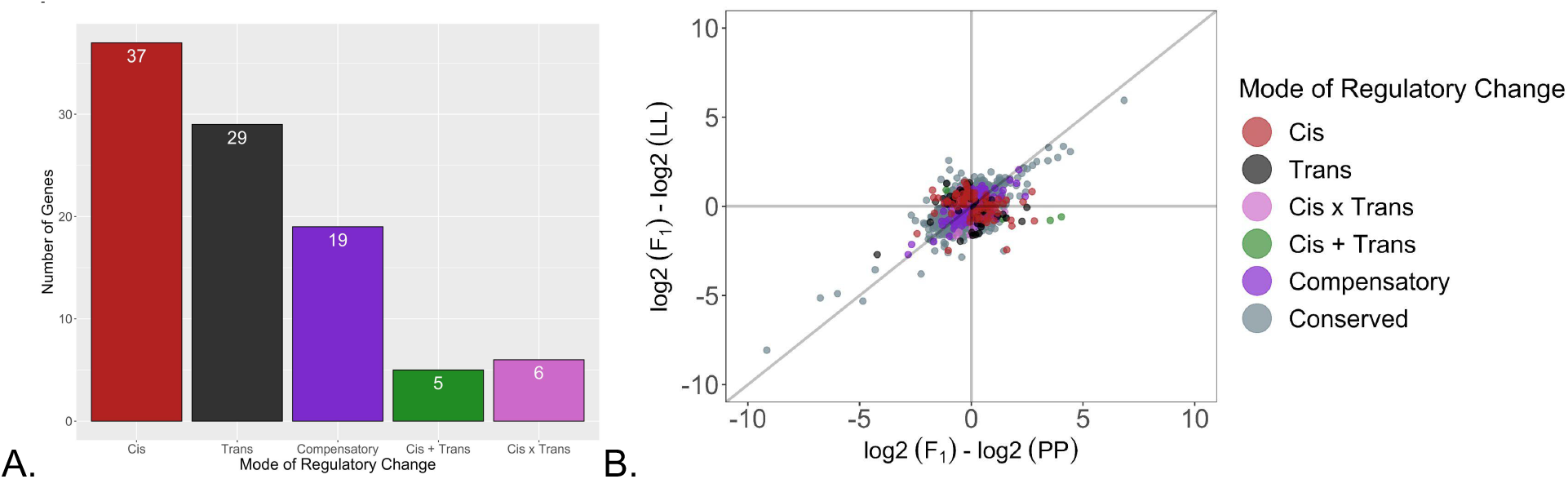
Regulatory architecture of differential expression in F_1_ oocytes. Both sets of F_1_s are analyzed together. Separate analysis of PL and LP showed similar results. (A) The number of genes in each category. (B) The distribution of genes from each category that are differentially expressed in the LL or PP direction.

## DISCUSSION

We investigate the changes in oocyte mRNA provisioning that accompany a common life-history transition in many animal taxa. *S. benedicti* has a developmental dimorphism directly associated with the classic life-history trade-off between egg size (offspring quality) and egg number (fecundity). Using this model system, we can investigate the regulatory architecture of this change in life-history with F_1_ crosses. Females of the two developmental modes produce eggs with a large difference in volume, and while F_1_ females produce intermediate egg sizes, F_2_ females produce a variable range of egg sizes, meaning that egg size is a quantitative trait^(30)^. Evolutionary theory predicts that the switch from planktotrophic to lecithotrophic larvae initially involves an increase in egg size followed by adaptive changes to larval morphology^(17,47–50)^. Despite the clear difference in egg size in *S. benedicti*, it is not known whether the larger lecithotrophic egg has quantitative differences in maternal mRNA provisioning, or if there are qualitative changes in maternal mRNAs in the egg that may affect ontogeny or larval phenotypes.

### Comparison of LL and PP egg gene expression

To investigate quantitative and qualitative changes in egg provisioning, we first compared the mRNA expression between eggs of planktotrophic and lecithotrophic mothers. We find that 43% (11,161) of all genes (26,216) in the *S. benedicti* genome are expressed in eggs of both morphs. This is more than in humans (36%^(51)^), *Xenopus laevis* (20%), and coral *Montipora capitata* (12%^(52)^) but fewer than some vertebrates: mice (63%), cows (44.7%^(53)^), zebrafish (49%^(54)^); and invertebrates: tunicate *Oikopleura dioica (63%*^(55)^) *Drosophila* (65%^(56)^). These studies represent a range of mRNA quantification techniques that likely lead to different estimates; however there is some conservation of genes that are maternally derived across all bilaterians^(57)^.

Of the 11,161 genes expressed in *S. benedicti* eggs, 10.2% are differentially expressed between the PP and LL eggs, which is a large difference considering it is due to allelic variation within a single species. Variation in mRNA provisioning can certainly occur within a species, particularly due to maternal condition or genotype, but this variation is usually small and pertains to a few genes. In *Drosophila melanogaster* the difference in oocyte gene expression between extreme environmental maternal conditions (starved mothers fed 5% of control mothers’ diet) is only ∼1.8%^(58,59)^. For mice, the difference in mRNA expression (of at least 4-fold) between pubertal (3 week old) and climacteric (58 weeks old) oocytes is 2%^(60)^. By comparison, in *Heliocidaris* urchins, planktotrophic and lecithotrophic larvae originate from different species, which are five million years diverged. Through early development ∼20% of their genes have different expression patterns, although this is not limited to the egg itself^(61)^.

Our findings show that the mRNA provisioning to eggs in *S. benedicti* is not simply caused by an increase in egg size with a modified amount of the same maternal mRNAs. A similar quantitative change in mRNA provisioning has been seen with egg increases in echinoderms^(61,62)^, and may indicate a general phenomenon associated with increased egg size. We find 18 genes are exclusively expressed in eggs of one morph. While not a particularly large subset, this is a substantial difference compared to most intraspecific analyses of differential expression. These exclusive genes highlight the possibility that the maternally controlled initiation of development is functionally distinct between the morphs. These genes should be the target of future studies investigating maternally directed development in this species.

Functional gene annotations reveal gene categories that are highly differentially expressed between PP and LL. Zinc finger proteins account for 12% (144) of differentially expressed genes and often serve transcriptional regulators. Because of poor annotation and lack of spiralian-specific genes in most commonly used gene databases, it is possible that such proteins may be lineage specific transcription factors, but this requires further functional validation. We also find that many known transcription factors are differentially expressed between PP and LL oocytes, including forkhead box protein (Sbene_G06980), polycomb protein (Sbene_G10227), kruppel-like factor 15 (Sbene_G10036), visual system homeobox 2 (Sbene_G05917), and paired-box protein (Sbene_G17489). Additionally, we identify genes involved in the Wnt signaling pathway, such as frizzled-5 (Sbene_G05398) and six putative copies of Notch (Sbene_G06517, Sbene_G09676, Sbene_G09677, Sbene_G13608, Sbene_G15020, Sbene_G18208). (Additional annotations in SFig. 1). Given that these genes are known actors in early development, it is possible that they could alter embryonic processes. This demonstrates the power of within-species hybrid transcriptomes to reveal candidates for subsequent functional studies.

### Comparison of reciprocal F_1_ gene expression with LL and PP

Because F_1_ females are heterozygotes (PL or LP), and they produce intermediate sized eggs, we may expect to see additive inheritance resulting in intermediate expression levels. However, few genes have an additive mode of inheritance and dominance is more pervasive. Because the within-group variance of the F_1_’s egg samples is larger than the variance between PP and LL, and because F_1_ expression is often intermediate, there may be statistical power issues that limit our ability to infer additivity (Fig. 3a). (In this case, these genes would be classified as ‘ambiguous’). An alternative possibility is that the F_1_’s eggs would contain the same gene expression as one of their parents due to dominance. Dominance early in development may indicate that the genes of one morph are required before the maternal-zygotic transition. A third (38.9%) of differentially expressed genes are dominant, and there is almost equal dominance between P and L alleles with a slight increase in L-dominance. We also see low levels of misexpression (over or underdominance: 0.8%, Fig. 2b) in the F_1_’s eggs, which is consistent with misexpression scaling with evolutionary divergence of the parents^(38)^.

Reciprocal crosses can also disentangle the contribution of each parent’s alleles to offspring phenotype^(63–66)^. Because we can make reciprocal crosses in *S. benedicti*, we are able to determine genes whose expression changes because of the parental background type alone: these are genes that have differential expression between LP and PL (Fig. 1). This analysis reveals the extent of differential expression and the particular genes that are impacted by genetic differences in the maternal background, which is an analysis that cannot be done with a unidirectional cross. If parent-of-origin effects are quantified from reciprocal crosses, they are reported during offspring development and have not been investigated during oogenesis. By comparing expression between the eggs of PL and LP females, we find that 41 genes exhibit parent-of-origin effects, which act in both the maternal and paternal direction. These expression differences occur when the F_1_ mother is provisioning her eggs, which means the transcript number an egg receives is due to the F_1_ mother’s parental cross direction. F_1_ females whose mother was PP provision their eggs differently than F_1_ females whose mother was LL, despite both F_1_ females having the same heterozygous genotype. (Although it is also possible that a minor amount of differentiation between F_1_ gene expression could be due to differences in the maternal mitochondrial haplotypes). Our results indicate that these parental-background effects persist and alter mRNA provisioning to eggs that make the next generation (an intergenerational effect on egg mRNA provisioning). To our knowledge, this is the first report of an intergenerational effect on offspring mRNA provisioning.

How do these parent-of-origin expression effects get passed to the next generation? Epigenetic modifications by maternally deposited mRNAs have been shown in vertebrates (zebrafish^(67)^, mouse^(68)^, *C. elegans*^(69,70)^). For invertebrates, most studies to date have identified sncRNAs as the maternally inherited drivers of epigenetic change^(71–74)^, although it is unclear if this is a pervasive mechanism across invertebrate taxa. There is evidence for epigenetic modifications via histone variants in some species^(75–77)^. Perhaps the simplest mechanistic explanation would be gene regulation by methylation, but studies on invertebrates suggest methylation is unlikely to be modifying expression^(78–80)^. It is possible that, while these genes have significantly different expression patterns, they have no functional effect on development. This role of these parent-of-origin genes in development should be investigated in future studies. However, maternal and paternal effects are evident for some larval phenotypes in *S. benedicti*^(30)^, suggesting that these maternally expressed differences could affect later developmental phenotypes. While the mechanism that causes parent-of-origin effects remains unknown, our study suggests there is some kind of epigenetic regulation of reproduction.

### Mode of Regulatory change

An advantage of our cross design is that we can determine the mode of regulation for differentially expressed genes. Typically regulatory analyses of *cis* and *trans* acting factors are conducted in hybrids^(35–38,81,59)^, however, we were able to carry out these analyses in intraspecific comparisons with the caveat that there are far fewer distinguishing polymorphisms between the alleles of the two parents. We focus on the genes that are differentially expressed between the two morphs. Therefore, we are not capturing the genetic architecture of genome-wide divergence between morphs, but rather the mode of inheritance in genes that are differently expressed. Nonetheless, we are able to identify the regulatory mode of 6.7% of the differentially expressed genes.

It is possible that differences in embryonic gene expression are due to a small number of *trans*-acting factors that can act pleiotropically to change expression of multiple genes. This is a parsimonious explanation for multiple phenotypic differences arising from a small number of changes to the genome. In closely related species, modifications to *trans* regulatory elements are the main driver of expression divergence^(81)^. More distantly related species have more *cis*-acting regulatory differences, that presumably are selected for and individually modified over evolutionary time^(35–38)^. We find that expression differences in F_1_ oocytes are explained by an almost even combination of both *cis*-acting and *trans*-acting factors, and a few instances of interactions (*cis* x/+ *trans*). For genes that are not differntially expressed between the two morphs, we do detect some compensatory changes, where the total expression is the same across parents because *trans-*acting factors are compensating for *cis-*acting changes or vice versa, and therefore expression levels differ in the F_1_s. Theory predicts that this occurs under stabilizing selection for expression level^(82)^, but over evolutionary time could lead to higher instances of misexpression. Our result demonstrates a complex genetic architecture in which maternal expression differences between the two morphs is due to both *trans* and *cis*-acting regulatory changes. Interestingly, this implies that numerous *cis* regulatory changes have evolved between these intraspecific types, in addition to *trans* acting factors.

We find that in *S. benedicti* maternal mRNA provisioning to eggs can vary significantly depending on the parents’ genotype. Differences in maternally provided expression in eggs can persist to the next generation, affecting how a mother packages mRNA to her own eggs. When comparing PP and LL eggs, we find significant expression differences for ∼10% of the genes, and this difference is only due to allelic variation across mothers. While changes in expression level make up the majority of the differences we see between morphs, there are also genes that exclusively appear in only one morph. (Although they are usually expressed in the F_1_s: SFigure 2). How these differences in mRNA egg provisioning affect downstream development remains to be determined, but this study shows that parental genetic effects on larval development could originate from morph-specific maternal provisioning.

## METHODS

### Animal Collection

We used lab-reared females descendent from two populations: planktotrophic worms are from Newark Bay Bayonne, New Jersey and lecithotrophic worms are from Long Beach, California. These populations are consistent with those used in previous genetic studies^(16,29,83,30)^. Planktotrophic and lecithotrophic individuals were reciprocally crossed to make F_1_ offspring PL and LP (maternal allele is listed first; Fig 1). Female F_1_s were reared until gravid, when we extracted and measured their eggs.

### Oocyte RNA collection and Library Prep

Eggs were dissected from female bodies by mechanical isolation of tissues in ice cold PBS. Extracted oocytes were counted and moved immediately to Arcturus PicoPure (Ref: 12204-01) RNA extraction buffer. The entire clutch of oocytes for a single-female was combined as one pooled sample. Libraries were constructed with the NEB UltraII Stranded RNA library prep kit for Illumina. Libraries were sequenced on two lanes of 150bp on the Illumina NovaSeq resulting in 80 million reads/library.

### Read quality trimming and Alignment

We use a combination of TrimGalore (cutadapt)^(84)^ and FastP^(85)^ to trim ambiguous bases. To remove rRNA we used SortMeRNA^(86)^, which identifies reads that map to a database of common eukaryotic rRNA sequences as well as annotated *S. benedicti* rRNA sequences. We aligned reads to the *S. benedicti* reference genome using HISAT2^(87)^ with strandedness, splice-site, and exon annotation guidance enabled using the default scoring parameters. SAM files were sorted by order of name using samtools sort, and feature-counting was done with HTSeq-count using the genome annotation file^(83)^. As the reference genome for *S. benedicti* is the planktotrophic morph, the final feature counts were averaged within each sample group to assess whether there was a significant bias towards this morph. We found planktotrophic samples had on average only 4% more of their total sequenced reads assigned to features than lecithotrophs. This is a small difference which demonstrates no significant mapping bias between the morphs. (STable 1)

### Normalization of F_1_ reads

When the F_1_ comparisons are incorporated in the analyses, all read counts were normalized together with the additional use of RUVg (RUVseq^(88)^) where a set of *a priori* housekeeping genes is used to standardize across samples (STable 2). This was necessary as the F_1_ samples were sequenced separately from the planktotrophic and lecithotrophic samples. A normalization step based on housekeeping genes enables us to compare between the parental and F_1_ sample groups and remove variance added by batch effects from the two separate sequencing runs. This does not introduce bias due to egg size differences because the mean egg size of the parental samples is equal to the mean egg size of F_1_ samples, and because the selected housekeeping genes do not exhibit differential expression between the planktotrophic and lecithotrophic eggs. This indicates that the expression of these housekeeping transcripts does not scale with egg size relative to other genes, making them a good reference for batch effects. After accounting for batch effects, variance stabilizing transformation was applied to counts in all four groups to make the PCA clustering, which is plotted for the first two principal components with ggplot2 (Figure 2A) and a heatmap of the expression values using the R package ‘pheatmap’ (Figure 2B).

### Differential Expression

Feature counts from all samples were concatenated and normalized together with DESeq2’s median of ratios method. This second sample normalization accounts for sequencing depth variance between each sample, independent of the sequencing batch. (Whereas the batch normalization step with RUVg permits comparisons between batches, the median of ratios normalization permits comparisons among all samples.) As part of the standard DESeq2 pipeline, p-values resulting from Wald tests were additionally corrected based on the Benjamini-Hochberg false discovery rate (FDR) algorithm to reduce the incidence of false positives for differential expression. Thresholds for significant differences were set as an FDR adjusted p-value of 0.05 or less, and an absolute fold change of more than 2x. Genes were considered very lowly expressed for a group if the group-mean normalized read count for a gene was below 10. If the other group’s mean normalized read count for those genes was above 150, then those genes were considered qualitatively differentially expressed. The significance level was adjusted for the analyses that include F_1_ oocyte RNAseq samples to accept FDR adjusted p-values of less than 0.10 in order to compensate for increased variability among the F_1_ samples.

### Mode of inheritance (Allele-specific effects)

To investigate the genetic basis of the expression differences found between PP and LL eggs, we made reciprocal F_1_ crosses (Fig. 1) and compared expression in their eggs back to PP and LL eggs. We assigned the primary mode of inheritance for each expressed gene using differential expression tests from DESeq2 according to established criteria (see STable 4)^(38,81)^. First, genes with parent-of-origin specific expression (differences in expression between F_1_s: PL and LP) were removed. The remaining genes were classified as being additive, dominant for one haplotype, or mis-expressed in heterozygotes (over/under-dominant). For example, a P-dominant gene is expressed at the same level in PP, PL and LP eggs, but differentially expressed in LL eggs (FDR adjusted p-value 0.1). Genes whose expression level is intermediate in the F_1_s (PL or LP) compared to PP and LL eggs are classified as additive, and genes expressed in F_1_s (PL or LP) at lower or higher levels than both PP and LL eggs are classified as underdominant and overdominant respectively.

### Parent of origin effects

For those genes exhibiting differential expression between reciprocal F_1_s, the same classification was applied to each group independently, and the results were compared. In these cases, genes are considered “maternal dominant” if they were classified as “P-dominant” in PL eggs, but not in LP eggs, or “L-dominant” in LP eggs but not in PL eggs. Conversely, genes are considered “paternal dominant” if they were classified as “L-dominant” in PL eggs, but not in LP eggs, or “P-dominant” in LP eggs but not in PL eggs.

### Mode of regulatory change

Alleles from F_1_’s eggs were assigned to one of the two parents by identifying fixed SNPs within the transcripts. We used HyLiTE^(89)^ (default parameters) to find SNPs and assign reads to the planktotrophic or lecithotrophic allele. To improve allele assignment rates, we included an additional 200 bp upstream and downstream of the input gene models to capture any reads which may align to untranslated regions not included in the annotations of coding sequences^(81)^.

The per-gene categorization of regulatory mode followed established empirical methods^(35,36,38,81)^. Only genes for which more than 20%, and no less than 10 total, of reads in the F_1_ samples could be assigned to a parent were considered. Three comparisons of each gene’s expression were made: First, we use genes that are differentially expressed between the parental morphs (PP and LL). Second, we calculate the allele-specific expression of each gene in the F_1_ (either PL or LP), using a negative binomial generalized linear model and Wald statistical tests using DESeq2 (v1.32.0). Third, we use a ratio of the differential expression of PP:LL alleles to the differential expression of LP:PL alleles. We use DESeq2, using the design (∼ W_1 + Geno * Ori) where W_1 is the normalization factor returned by RUVg, Geno identifies reads as either a P or L allele, and Ori identifies the reads as originating from the parentals or F_1_ samples. Based on the results of the three above comparisons, DE genes were categorized as either in “cis”, “trans”, “cis + trans”, or “cis * trans.”^(35,36,38,81)^. (Further explanation of cis/trans categories in STable 3).

### Functional Annotations

Previously established genome annotations^(83)^ were improved by performing a BLASTx search against the complete UniProt/SwissProt protein databases, adding any annotations that had an e-value less than 10^−30^. Gene functional information was retrieved from the UniProtKB database.

We used gene ontology (GO) information (SFig. 1), but did not test for term enrichment because we could not assign terms to the majority of the genes in our dataset.

## DATA ACCESSIBILITY

All data and analyses will be made publicly available. Raw reads will be submitted to NCBI’s short read archive. Updated functional annotations are submitted to NCBI. All analyses and datasets are available in the supplemental files, which contain input data and a Rmarkdown file.

## ACKNOWLEDGEMENTS

We would like to thank S. Cole for assistance with animal care. Thanks to members of the C. Zakas lab and G. Wray lab (2019-2022) for feedback on the project. Thanks to M. Rockman, G. Wray, and H. Devens for specific feedback on the manuscript.

## AUTHOR CONTRIBUTIONS

N. D. Harry is responsible for data curation and formal analysis. C. Zakas is responsible for conceptualization, funding acquisition, and project administration. Authors shared responsibility for methodology, investigation, visualization, and writing and reviewing the manuscript.

## SUPPORTING INFORMATION CAPTIONS

### Tables

*Supplement Table 1. Sequencing and read processing results for all libraries*.

*Supplement Table 2: Selected Housekeeping genes (for RUVg)*

*Supplement Table 3: Criterion for regulatory mode assignments*

*Supplement Table 4: Criterion for mode of inheritance assignments*

*Supplement Table 5: Annotation of transgenerational differentially expressed genes*

*Supplement Table 6. Genes exclusive to one group: mean counts and annotations*

*Supplement Table 7. F*_*1*_ *egg expression patterns for exclusive genes*.

### Figures

S*upplement Figure 1: REVIGO tree-map of GO terms associated with differentially expressed genes between PP and LL eggs*

*Supplement Figure 2. Differences in F*_*1*_ *egg expression for genes exclusive to LL or PP eggs. Supplement Figure 3. Shared gene expression across all groups*.

### Files

*<u>Supplement input data and R files</u>*

